# Cave pools in Carlsbad Caverns National Park contain diverse bacteriophage communities and novel viral sequences

**DOI:** 10.1101/2024.09.12.612726

**Authors:** Joseph Ulbrich, Nathaniel E. Jobe, Daniel S. Jones, Thomas L. Kieft

## Abstract

Viruses are the most abundant biological entities on Earth, and they play a critical role in the environment and biosphere where they regulate microbial populations and contribute to nutrient cycling. Environmental viruses have been the most studied in the ocean, but viral investigations have now spread to other environments. Here, viral communities were characterized in four cave pools in Carlsbad Caverns National Park to test the hypotheses that (i) viral abundance is ten-fold higher than prokaryotic cell abundance in cavern pools, (ii) cavern pools contain novel viral sequences, and (iii) viral communities in pools from developed portions of the cave are distinct from those of pools in undeveloped parts of the same cave. The relationship between viral and microbial abundance was determined through direct epifluorescence microscopy counts. Viral metagenomes were constructed to examine viral diversity among pools, identify novel viruses, and characterize auxiliary metabolic genes (AMGs). Bacterial communities were characterized by 16S rRNA gene amplicon sequencing. Epifluorescence microscopy showed that the ratio of viral-like particles (VLPs) to microorganisms was approximately 22:1 across all sites. Viral communities from pools with higher tourist traffic were more similar to each other than to those from less visited pools, although surprisingly, viruses did not follow the same pattern as bacterial communities, which reflected pool geochemistry. Bacterial hosts predicted from viral sequences using iPHoP showed overlap with both rare and abundant genera and families in the 16S rRNA gene dataset. Gene-sharing network analysis revealed high viral diversity compared to a reference viral database as well as to other aquatic environments. AMG presence showed variation in metabolic potential among the four pools. Overall, Carlsbad Cavern harbors novel viruses with substantial diversity among pools within the same system, indicating that caves are likely an important repository for unexplored viromes.

## Introduction

Cave systems have fascinated explorers and scientists with their isolated environments and unique ecology. Caves are natural voids within the subsurface of the Earth that are large enough for human exploration [1]. They typically feature isolated ecosystems with low input from outside influences such as nutrients and light that serve as invaluable resources for research. These extreme ecosystems are home to specially adapted organisms, across all three evolutionary domains, that provide insights into extraterrestrial life and the evolution of life on Earth [2, 3]. Carlsbad Caverns National Park is home to North America’s largest publicly accessible cave, Carlsbad Cavern, the largest chamber of which is called the Big Room. Carlsbad Cavern has approximately 500,000 tourist visits annually (https://www.nps.gov/cave/learn/news/interesting-facts-about-carlsbad-caverns.htm) and has also been the subject of various scientific studies. Microbial surveys have been conducted there [2–6], but investigation of viruses has been limited. Previous viral research at Carlsbad Cavern focused primarily on viruses in bat populations [7, 8]. Environmental viruses have received very little attention in caves. A recent viral search was conducted in a cave-like environment: an anchialine system, specifically a near-shore underground estuary in karst terrain accessed via a sinkhole [9]; the authors reported abundant, diverse bacteriophages and archaeal viruses containing auxiliary metabolic genes encoding sulfur- and phosphorous-related functions.

Viruses, along with their hosts, are known to inhabit a wide variety of environments, including those that are extreme with regard to temperature, pH, pressure, or salinity [10, 11]. Although less researched than viruses that infect humans, environmental viruses that infect bacteria (bacteriophages) and archaea are vital to the health of ecosystems and are drivers of evolution [10, 12, 13]. Carbon cycling in oceans and food web stability have been directly influenced by viruses through the viral shunt, which is the release of nutrients as a result of host cell lysis [14, 15]. It is estimated that viruses lyse 20% of bacteria in the surface ocean daily [15] and that 6.8-42% of marine bacterial organic carbon flows through the viral shunt [14]. Additionally, as a result of host cell lysis, viruses contribute to microbial evolution by selecting virus-resistant microbes [16]. Viruses also play a major role in lateral and vertical gene flow between organisms as a result of their lytic and lysogenic mechanisms [12, 16]. This gene flow can have ecological implication, such as by transferring genes for photosynthetic machinery [17] or triggering cell death [18]. Viruses within cavern systems could similarly impact geochemical dynamics and microbial evolution in Earth’s subsurface.

Traditionally, environmental phage studies have been culture-dependent, requiring the extraction of viral-like particles (VLPs) from the environment, followed by the propagation of phages in host cells [19]. This approach is severely limiting due to the inability to cultivate the majority of microorganisms in the lab [20, 21], the difficulty of identifying and isolating phages, the lack of plaque formation, and pseudolysogeny [19, 22]. Most bacteria and archaea are unculturable “microbial dark matter” [20, 21] and they host a corresponding “viral dark matter” [23] that eludes culture-dependent analysis. Culture-independent approaches based on DNA sequencing circumvent this culture bias. The majority of phages and archaeal viruses are thought to be DNA viruses [24], which simplifies analyses, although RNA phages are also known [25]. There are no universally conserved viral sequences, so universal primers for PCR amplification are not possible, but high-throughput metagenomic sequencing can be used to characterize environmental viromes. Direct sequencing and analysis of viral genetic material involves (i) viral sample recovery, such as by tangential flow filtration or sorption to iron oxide [26], (ii) VLP purification, (iii) VLP concentration, and (iv) genomic sequencing [19].

The goal of this study was to investigate viral communities from cavern water pools in Carlsbad Caverns National Park. The following hypotheses were tested: i) viral abundance is ten-fold higher than prokaryotic cell abundance in cavern pools, (ii) cavern pools contain novel viral sequences, and (iii) viral communities in pools from developed portions of a cave are distinct from those of pools in undeveloped parts of the same cave. Further investigation was performed to determine the functional genes and role they play in the environment, and explore the relationship between DNA viruses and microorganisms present in the sample environments.

## Material and Methods

### Study Site

This study was conducted at Carlsbad Caverns National Park, NM. Sampling sites within Carlsbad Cavern (Fig. 1) were selected based on their relative amounts of human traffic and their size. Half of the samples were taken from open access self-guided areas in the artificially lighted Green Lake Room and Big Room adjacent to the paved tourist route. Green Pool is relatively broad (3 × 6 m) and shallow (1 m depth); Longfellow’s Bathtub is long (13 m), narrow (1.5) and deep (2 m). Both of these illuminated pools have visible accumulations of photosynthetic biofilms, i.e. “lampenflora” [5]. The remaining samples were collected from pools in unlighted areas that are visited only via scheduled small-group tours in Lower Cave and Left-Hand Tunnel. Surface dimensions of Lower Cave Pool are ∼6 × 2.5 m and the depth is ∼1.8 m. Iron Pool in the Left-Hand Tunnel has similar dimensions, and while it is in an area that is remote from the cave entrance, it is along a flyway to a major bat colony and had fresh bat guano pellets on the bottom of the pool.

**Fig. 1.**
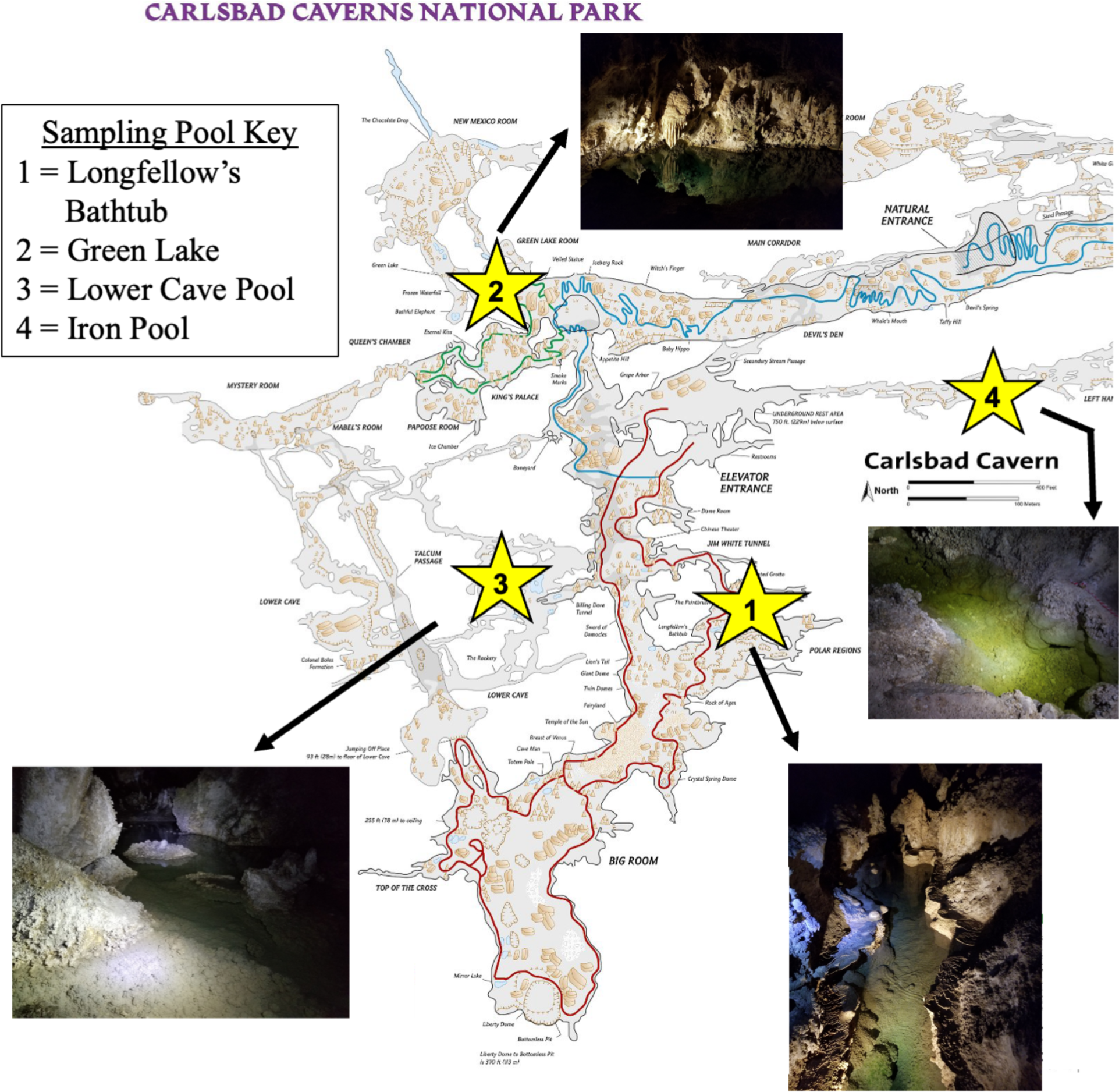
Map of Carlsbad Cavern showing the locations of the four pools sampled in this study.

### Physical and Chemical Measurements

Temperature and pH of cave pool water were measured with a Hanna Instruments pHep4 pH meter. Water samples (250 mL for analysis of general water chemistry and 125 mL for trace element analysis) were collected from each pool. Specific conductance was measured using the Environmental Protection Agency (EPA) method 120.1. pH was measured using the EPA method 150.1. Cations were measured using inductively coupled plasma optical emission spectroscopy (ICP-OES) using EPA method 200.7. Anions were measured using ICP-OES under EPA 300.0. The alkalinity test was performed under EPA 310.1. Trace metals were measured via inductively coupled plasma mass spectrometry (ICP-MS) under EPA 200.8. Hardness was calculated via the SM 2340B method (http://standardmethods.org). All chemistry and trace element measurements were performed at the New Mexico Bureau of Geology and Mineral Resources.

### Biological Sampling

The overall scheme for biological processing and analyses is shown in Fig. 2. Filtered and unfiltered samples were also used for quantifying VLPs and cells, respectively. Pool water (10 L) was filtered on-site, from which the filtrate was used for viral sequence analysis and the filters were used for bacterial 16S rRNA gene amplicon sequencing.

**Fig. 2.**
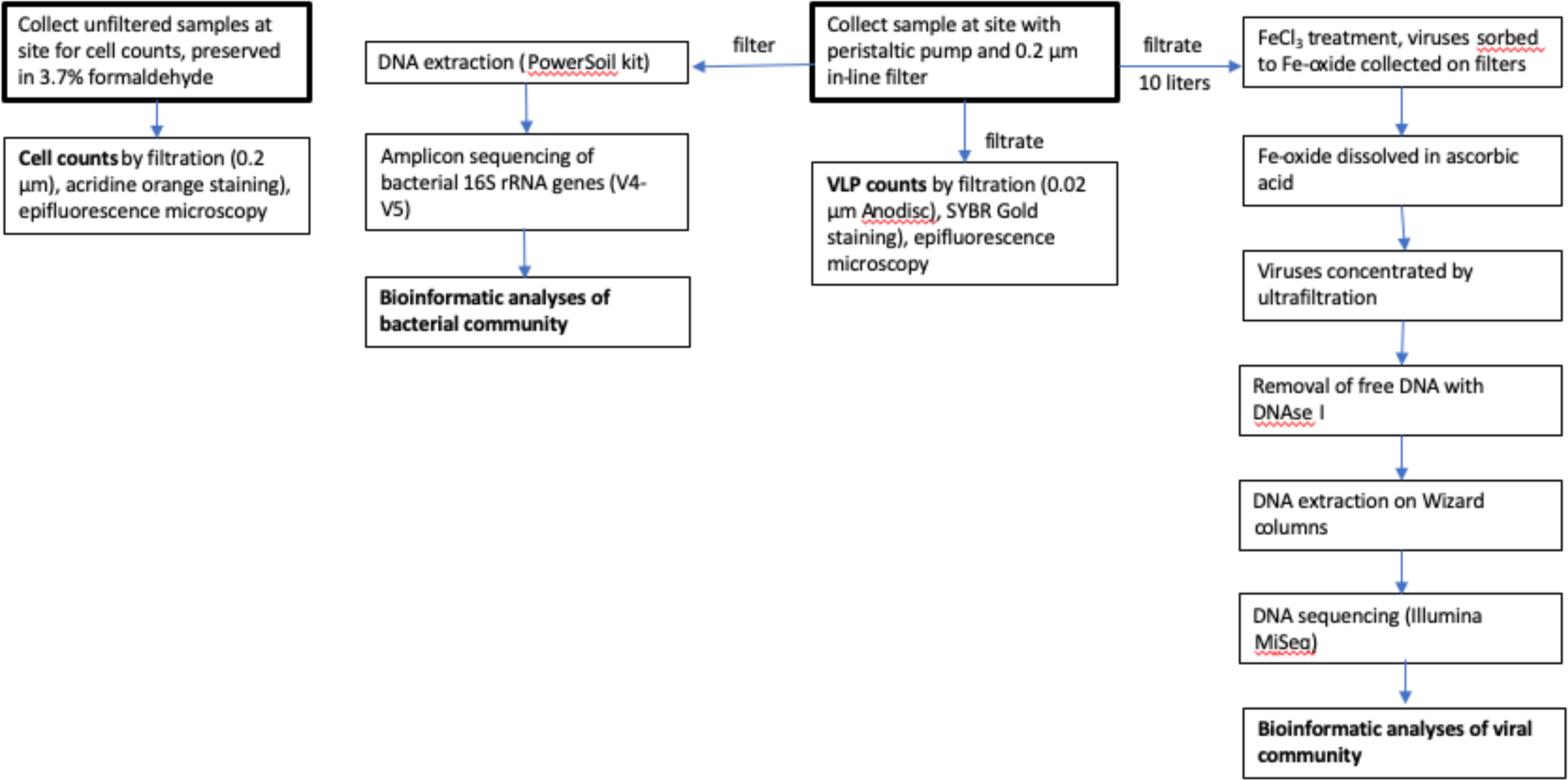
Flow chart diagramming biological sample collection and processing

### Quantifying Cells and VLPs

Two 40-mL samples were collected for microbial quantification at each site and preserved with formaldehyde (3.7%). Viral quantification samples were collected using a 0.2-µm inline filter apparatus, collecting two 45-mL samples of filtrate from each pool. All samples were collected into sterile 50-mL centrifuge tubes and placed on ice for transportation. All samples were refrigerated at 4 °C to prevent cell lysis until microscopic counts were performed.

Cell counts were performed using acridine orange staining and epifluorescence microscopy [27]. Unfiltered pool samples (10 mL each) were vacuum-filtered through polycarbonate filters (25-mm diameter, 0.2-µm pore size, black) backed by a nitrocellulose filter (25-mm diameter, 0.2-µm pore size, white). In the filter tower, 5 drops of acridine orange solution (1.0 g L^-1^, filter sterilized) were added and mixed with 5 mL of buffer solution (25 g NaHCO_3_, filter sterilized) and allowed to stand for three minutes. The staining solution was passed through the membrane and the filter was placed onto a microscope slide to dry in the dark. After ∼20 minutes, a coverslip was placed onto the filter with a drop of low-fluorescence immersion oil. Slides were kept in the dark until direct counting could be performed using epifluorescence microscopy at 630× magnification. Ten fields were counted for each of the fields and an average cell concentration was calculated for each cavern pool sampled.

VLP quantification was performed following the method of Noble and Fuhrman [28]. Viruses in the 0.2-µm filtrate were captured on a 0.02-µm Anodisk filter. SYBR Gold was used as a stain in place of SYBR Green I as a suitable alternative [29]. VLPs were counted using epifluorescence microscopy.

### Capture and Concentration of Viruses

Water samples (10 L each) were collected from each pool using a sampling apparatus consisting of a 0.2-µm Whatman 36AS in-line filter attached to a 2.4-m length of silicone tubing. Samples were drawn through the tubing using a Master Flex E/S Portable peristaltic pump and collected into autoclaved 10-L polypropylene carboys. Following collection of the filtered water, the Whatman 36AS filters were placed into Whirl-Pak bags and the water in the carboys was treated with 1 mL of 10 g L FeCl_3_ stock solution following the method of John et al. [26]. Water samples and filters were placed on ice and transported to New Mexico Tech, where the water samples were refrigerated at 4°C for further processing and the 0.2-µm Whatman filters were stored at −80°C for bacterial amplicon sequence analysis. The 10-L water samples containing iron oxide and sorbed viruses were then concentrated onto 0.8 µm pore size polycarbonate filters by vacuum filtration. Following filtration, sets of three 0.8 µm filters were placed into 50-ml centrifuge tubes and stored at 4 °C.

Iron oxide dissolution and viral resuspension were performed following the method of John et al. [26], modified for 10-L water samples. Three 0.8-µm polycarbonate filters with bound Fe-oxide and viruses from each pool were placed into 50-ml mL centrifuge tubes and 10 mL of buffer (0.25 M ascorbic acid, 0.2 M Mg(II)-EDTA, pH 6-7) was added to dissolve the iron oxide and resuspend the viruses and the tubes were incubated at 4°C with rotary mixing overnight.

Viral suspensions were concentrated using ultrafiltration devices and centrifugation. Samples were first reduced to ∼4 mL using Millipore Amicon centrifugal filter tubes (catalog # UFC910024). A 3-mL subsample of each sample was set aside while the rest of the viral suspension was added to the upper reservoir of the Amicon ultrafiltration tube. Several rounds of centrifuging at 1000 × g for 1 minute each at 4 °C in a swinging-bucket rotor were performed, discarding the flow through, until ∼1 mL remained in the upper reservoir for each sample. The 1 mL was removed, saved, and 1.5 mL (half) of the set aside subsample was added to the upper reservoir. The bottom of the upper reservoir was parafilmed and the Amicon device was vortexed on medium for 20 seconds. After vortexing, this process was repeated until 4 mL of viral suspension remained for each pool. The sample volumes were further reduced using Pall Nanosep centrifugal filter tubes (catalog # OD100C34) until a final volume of 405 µL was achieved. Centrifugation was carried out under the same parameters and resuspension rounds were performed with 10 µL instead of 1.5 mL. Centrifugation duration and number of cycles varied based on the volume of samples and the amount of material in each sample.

Concentrated viral suspensions were treated with DNase I to remove free DNA based on the method of Hurwitz et al. [30]. DNase stock solution of 40,000 U mL^-1^ was diluted with 10× reaction buffer (100 mM Tris-HCl, pH 7.6, 25 mM MgCl_2_, 5 mM CaCl_2_) to a working concentration of 1 U mL^-1^, and samples were treated with a 1:10 dilution of enzyme solution to viral suspension. Reactions were incubated for 2 hours at room temperature and DNase was then inactivated by adding EDTA (100 mM final concentration). This resulted in a final volume of 500 µL.

DNA was extracted from viral suspensions using Wizard Minicolumns (Promega). Each 500-µL viral sample was mixed with 1 mL DNA purification resin. The resin/viral suspensions were washed through Wizard Mini columns, then washed with 2 mL of 80% isopropanol, and centrifuged at 10,000 × g for 2 minutes. The residual liquid was removed and 100 µL 80°C molecular grade water was added to the top of the mini columns. The columns were vortexed gently for 10 seconds and allowed to stand for one minute. The mini columns were centrifuged at 10,000 × g for 30 seconds and DNA was eluted.

Extracted viral DNA was sequenced at the University of Minnesota Genomics Center using Illumina MiSeq sequencing. Ampure bead cleaning (Beckman-Coulter) was performed on all samples prior to sequencing, concentrating samples to 15 µL.

### Viral Sequence Analysis

Following library construction, low quality reads were filtered and removed using Sickle 1.33 [31], and residual sequencing adapters were removed from 3’ ends using Cutadapt 3.2 [32]. Assembly was performed on paired and unpaired reads using metaSPAdes [33], with reads from all four sites co-assembled. Contigs larger than 1.5 kilobase pairs (kbp) were recognized as viruses if they had a final VirSorter2 v2.2.4 [34] score of at least 0.9 and VirFinder v1.1 [35] score of at least 0.9 and a p-value less than 0.05. Viral contigs that met each criterion were taxonomically clustered using vConTACT2 v0.11.1 [36] by calling genes in the high confidence viral contigs using Prodigal v2.6.3 [37] and the -meta flag and clustering using ‘ProkaryoteViralRefSeq211-Merged’ database. Cluster networks were visualized using Cytoscape [38]. Additional clustering was done using the 4May2024 INPHARED database [39] and used in conjunction with graphanalyzer v1.6.0 [40] in order to help in cluster interpretation and taxonomic assignment of tightly clustered contigs. Host prediction was done using iPHoP v1.3.3 [41] with the Aug_2023_pub_rw database. Auxiliary metabolic genes in the viral contigs were predicted using VIBRANT v1.2.0 [42]. A crude classification of viruses was performed by BLASTing all viral contigs using BLASTX and all predicted proteins using BLASTP against the RefSeq virus database, and then assigning them to families or higher categories by using the least common ancestor (LCA) algorithm in MEGAN v. 6.25 to summarize the taxonomic assignment of BLAST matches within 10% of the bit score of the highest-scoring match. Viral genome annotation for complete circular genomes and high quality draft genomes as designated by VIBRANT was performed with pharokka v 1.7.3 using the –m flag for metagenomes and -s to make separate outputs for each contig (https://github.com/gbouras13/pharokka?tab=readme-ov-file#citation) and phold version 0.1.4 (https://github.com/gbouras13/phold) using the pharokka output with the -separate and -cpu flags.

### Bacterial 16S rRNA Gene Amplicon Sequence Analysis

DNA was extracted from Whatman 36AS filters using the MoBio (now Qiagen) PowerSoil DNA isolation kit (MoBio Laboratories Inc., Carlsbad, CA, USA). rRNA gene libraries were prepared by amplifying the V4 hypervariable region of the small-subunit rRNA with primers 515f/806r [43]: 515f modified, GTG YCA GCM GCC GCG GTA A; 806r modified, GGA CTA CNV GGG TWT CTA AT) and Nextera adaptors (forward tail, TCG TCG GCA GCG TCA GAT GTG TAT AAG AGA CAG; reverse tail, GTC TCG TGG GCT CGG AGA TGT GTA TAA GAG ACA G). PCR conditions were 5 min for initial denaturation at 94 ◦C, followed by either 30 or 35 cycles for 45 s at 94 ◦C, 60 s at 50 ◦C, and 90 s at 72 ◦C, using the HotStarTaq Plus polymerase kit (Qiagen). Libraries were barcoded, pooled, and sequenced on an Illumina MiSeq platform (paired end, 2 × 300 bp) at the University of Minnesota Genomics Center. We also attempted to prepare rRNA gene libraries for bacterial, archaeal, and eukaryote communities by submitting DNA extracts for rRNA gene amplicon sequencing using the full service library preparation option with primers for the V4, V9 (18S), and Arc 515F/Arc915R primer pair [44–46].

Raw fastq sequences were analyzed using a procedure similar to that of [47]. Forward reads were trimmed to an average quality of ≥28 with sickle (https://github.com/najoshi/sickle), reads with residual adaptors were removed with cutadapt [32], and then forward and reverse reads were assembled with PEAR [48]. Operational taxonomic unit (OTU) calling at 97% identity and chimera removal were performed with the UPARSE pipeline [49], and the taxonomy of representative sequences for each OTU was determined against the Silva database v.138 [50] using mothur [51].

Statistical analyses were performed in R v. 4.3.1 [52] using the vegan package v. 2.6-4 [53]. Raw OTU counts were standardized using the Hellinger transformation, and hierarchical agglomerative cluster analysis was computed using Bray-Curtis distance and UPGMA (unweighted pair group method with arithmetic mean) linkage. For two-way cluster analysis, libraries were clustered (R-mode clustering) based on the 50 most abundant OTUs. Clustering of sites based on geochemical parameters was performed using the parameters in Table 1, using the same clustering techniques after for rRNA amplicon libraries but with geochemical parameters standardized to the maximum measured value for each variable. Clustering of viral communities was performed using Bray-Curtis distance and UPGMA linkage, based on the coverage of each viral contig in each sample, after first converting viral coverage to proportion of the total coverage of all viral contigs in that sample. R-mode cluster analysis of viral communities was performed for putative viral contigs that were more 1% of the total coverage in at least one sample. Diversity indices were calculated using Microsoft Excel and standard formulas for calculating Shannon [54], Simpson’s [55], and Chao1 [56] indices.

**Table 1.** Summary of cave pool geochemistry.

| Parameter | Lower Cave Pool | Longfellow's Bathtub | Iron Pool | Green Lake |
| --- | --- | --- | --- | --- |
| Specific conductivity ($\mu\text{S cm}^{-1}$) | 373 | 982 | 14700 | 419 |
| pH | 8.2 | 7.8 | 8.6 | 8.2 |
| Alkalinity ($\text{meq L}^{-1}$ as $\text{CaCO}_3$) | 178 | 75.3 | 531 | 205 |
| Major anions ($\text{mg L}^{-1}$) | | | | |
| Bromide | 0 | 0 | 14.7 | 0 |
| Chloride | 4.42 | 12.4 | 1270 | 5.212 |
| Fluoride | 0.64 | 0.74 | 0 | 0.78 |
| Nitrate | 12.4 | 14 | 5030 | 10.3 |
| Nitrite | 0 | 0 | 5.05 | 0 |
| Phosphate | 0 | 0 | 0 | 0 |
| Sulfate | 15.8 | 428 | 3640 | 16.3 |
| Major cations ($\text{mg L}^{-1}$) | | | | |
| Ca | 18.7 | 138 | 28.8 | 20.1 |
| Fe | 0 | 0 | 0 | 0 |
| Mg | 36.6 | 48.2 | 1810 | 43.5 |
| K | 0.514 | 1.8 | 354 | 0.466 |
| Si | 9.27 | 18.4 | 1.04 | 9.29 |
| Na | 5.85 | 11.6 | 1200 | 6.59 |
| Sr | 0.258 | 0.213 | 0.147 | 0.114 |

## Results

### Environmental Conditions

Chemical parameters varied among the four pools (Table 1, Supplementary Table S1). The pH varied from 7.8 to 8.6, a range that is consistent with buffering by the carbonate cave bedrock. Total dissolved solids, along with alkalinity, conductivity, and the concentrations of major anions (chloride, nitrate, and sulfate) and cations (calcium, magnesium, potassium, and sodium) were highest in Iron Pool (Table 1). Nitrate was more than 400× higher in Iron Pool than in other pools, at 5030 mg L^-1^, consistent with the influence of bat guano at that site. Similarly, the ICP-MS analyses showed highest concentrations of boron, lithium, molybdenum, and selenium in Iron Pool (Supplementary Table S1). Longfellow’s Bathtub was saltier than either Lower Cave Pool or Green Lake, which both had conductivities less than 500 μS cm^-1^, but all three lakes were much fresher than Iron Pool.

### Viral and Bacterial Abundance

The abundances of VLPs and bacteria were relatively consistent among the four pools. VLPs ranged from 1.15 × 10^5^ to 4.15 × 10^5^ VLPs mL^-1^ and bacteria ranged from 7.00 × 10^3^ to 1.44 × 10^4^ cells mL^-1^ (Table 2). Iron Pool had the highest concentrations of both. Ratios of VLPs to bacteria ranged from 15:1 to 29:1, with a mean ratio of 22:1 (Table 2).

**Table 2.** Abundance of viral-like particles (VLPs) and bacterial cells in the four cavern pool samples.

| **Cavern Pool** | **VLPs mL<sup>-1</sup>** | **Bacteria mL<sup>-1</sup>** | **VLP:prokaryote**<br>**ratio** |
| --- | --- | --- | --- |
| Longfellow's Bathtub | 1.2 x 10<sup>5</sup> | 7.9 x 10<sup>3</sup> | 15 |
| Green Lake | 1.8 x 10<sup>5</sup> | 7.0 x 10<sup>3</sup> | 25 |
| Lower Cave Pool | 1.7 x 10<sup>5</sup> | 9.4 x 10<sup>3</sup> | 18 |
| Iron Pool | 4.2 x 10<sup>5</sup> | 1.4 x 10<sup>4</sup> | 29 |
| **Average** | **1.7 x 10<sup>5</sup>** | **8.1 x 10<sup>3</sup>** | **22** |

### Viral Community Analysis

Illumina MiSeq sequencing of DNA from the viral extracts from all four pools generated >8 × 10^7^ raw reads. Co-assembly of reads from all samples led to more than 3000 contigs longer than 1500 bp. Of these, VirSorter2 and VirFinder detected a total of 1505 contigs that can be confidently considered to represent viruses; based on the samples from which they are most abundant, these putative viruses are relatively evenly distributed among the four samples (Green Lake: 378 contigs, Iron Pool: 278 contigs, Longfellow’s bathtub: 549 contigs, Lower Cave Pool 300 contigs, Fig. 3). Vibrant identified 1243 of these 1505 contigs as bacteriophage sequences, but only 36 of these were assembled into complete circular viral genomes. 418 viral contigs clustered tightly with other viruses from the RefSeq viral database provided with vConTACT2. Many of these are novel viruses as shown by network analysis (Fig. 4). Using the INPHARED database, viral contigs that clustered tightly to the database viruses were categorized using graphanalyzer. Of the 1505 viral contigs, 53 were able to be classified to the Caudoviricetes class while 15 of these were classified to the family level. We also used BLAST to compare all predicted proteins from the 1505 viral contigs against the RefSeq viral database, which shows that 84% of proteins had no BLAST matches and 15% matched viral proteins in the database. According to the NCBI taxonomic classification, Siphoviridae, Myoviridae, and Podoviridae were the most abundant groups of viruses in the cavern pools (Fig. 5).

**Fig. 3.**
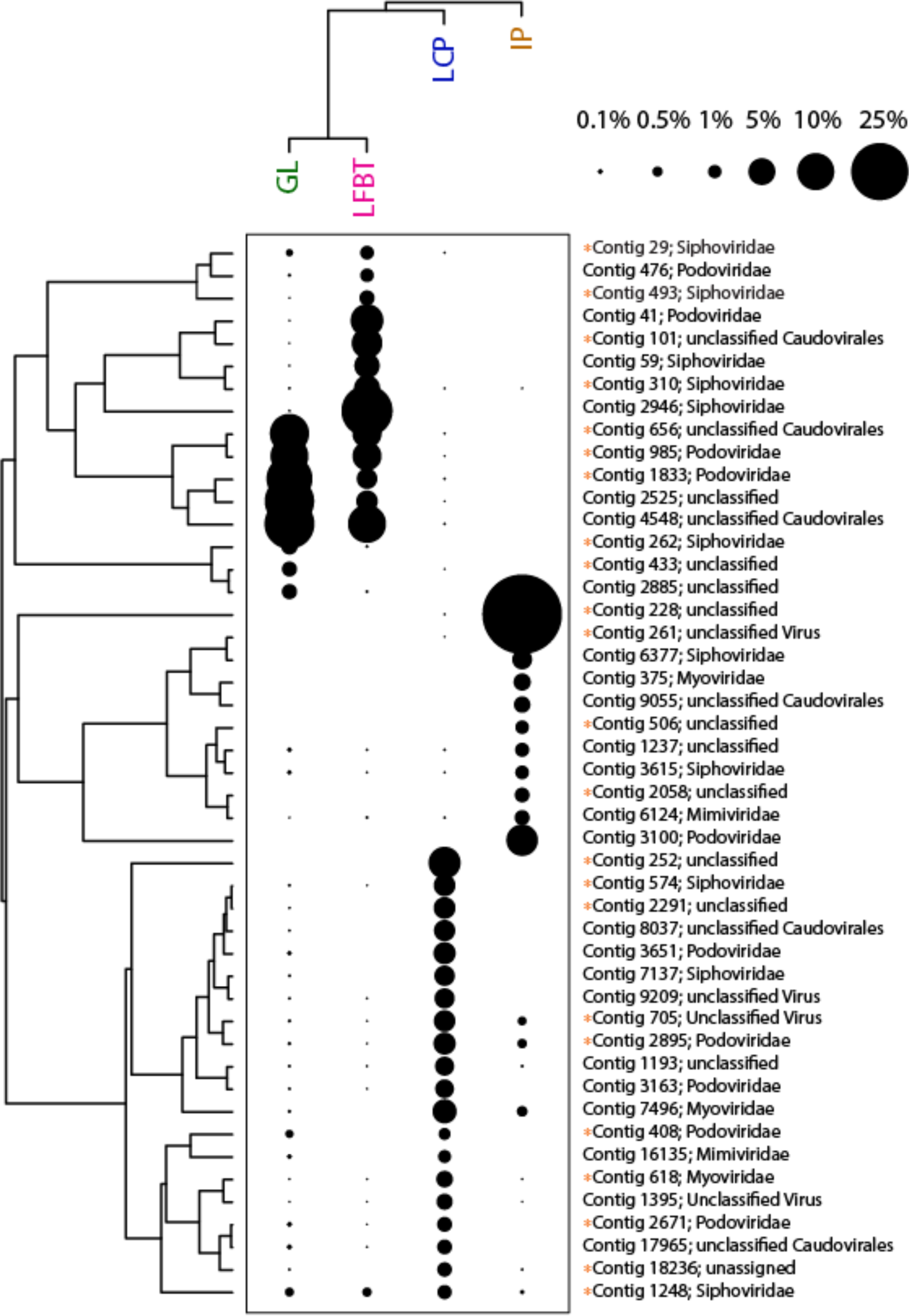
Two-way cluster analysis of metaviromes from the four cave pools, based on coverage of viral contigs in each metavirome library. The Q-mode cluster analysis was calculated with all 1505 putative viral contigs, while the R-mode cluster analysis was only those OTUs that were more than 1% relative abundance (as a percentage of total coverage of viral contigs in each sample). The sizes of the points scale with relative abundance. The classification of each contig was determined using the LCA algorithm in MEGAN, based on BLAST searching against the RefSeq viral database. Unclassified viruses are designated with an orange dot before the contig number.

**Fig. 4.**
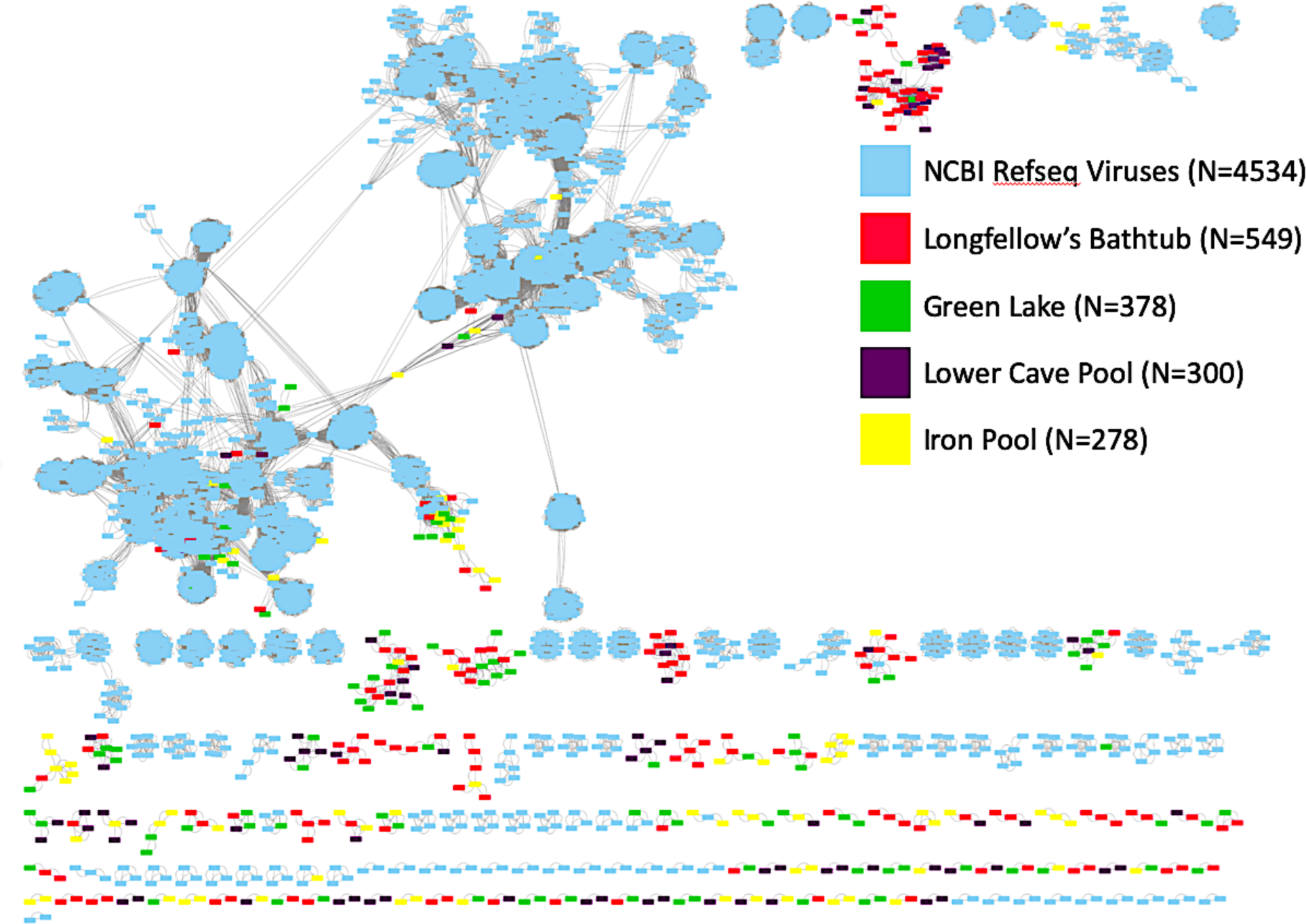
Network analysis. Light blue boxes represent known phages in the vConTACT2 database, green boxes are for Green Lake, orange boxes are for Iron Pool, the purple boxes are for Lower Cave Pool, and the red boxes are for Longfellow’s Bathtub. For phages that occurred in more than one pool, the pool with the highest coverage of that phage is indicated by the color for that pool. The blue boxes represent viral contigs that matched to NCBI database reference sequences.

**Fig. 5.**
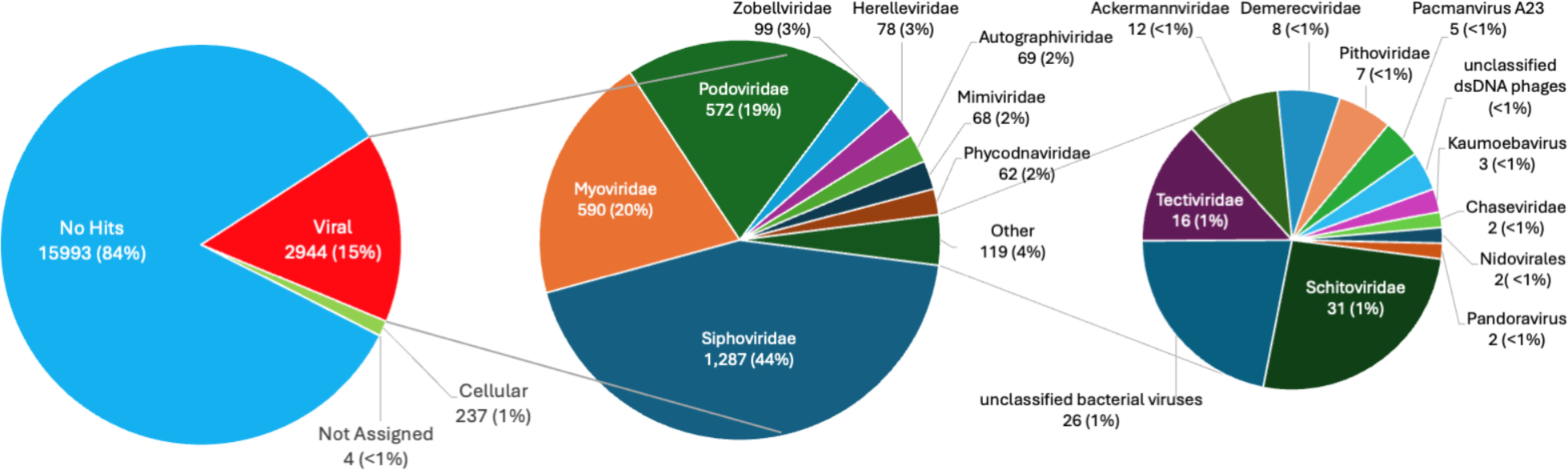
Viral DNA extract reads mapped using BLASTP and the LCA algorithm in Megan.

VIBRANT identified 71 different auxiliary metabolic genes (AMGs) among a total of 257 metabolic genes found in the four samples (Fig. 6). Of the four pools, Longfellow’s Bathtub had the majority of the AMGs mainly in the categories of carbohydrate metabolism (n = 44), metabolism of cofactors and vitamins (n = 33), and glycan biosynthesis and metabolism (n = 20). The most abundant categories for Green Lake and Lower Cave Pool were similar in that metabolism of cofactors and vitamins was the most abundant AMG category (Green Pool: 26, Lower Cave Pool: 20). Iron Pool had the fewest AMGs in the viral contigs most abundant in the pool with carbohydrate metabolism (n = 11) and amino acid metabolism (n = 8) being the most abundant.

**Fig. 6.**
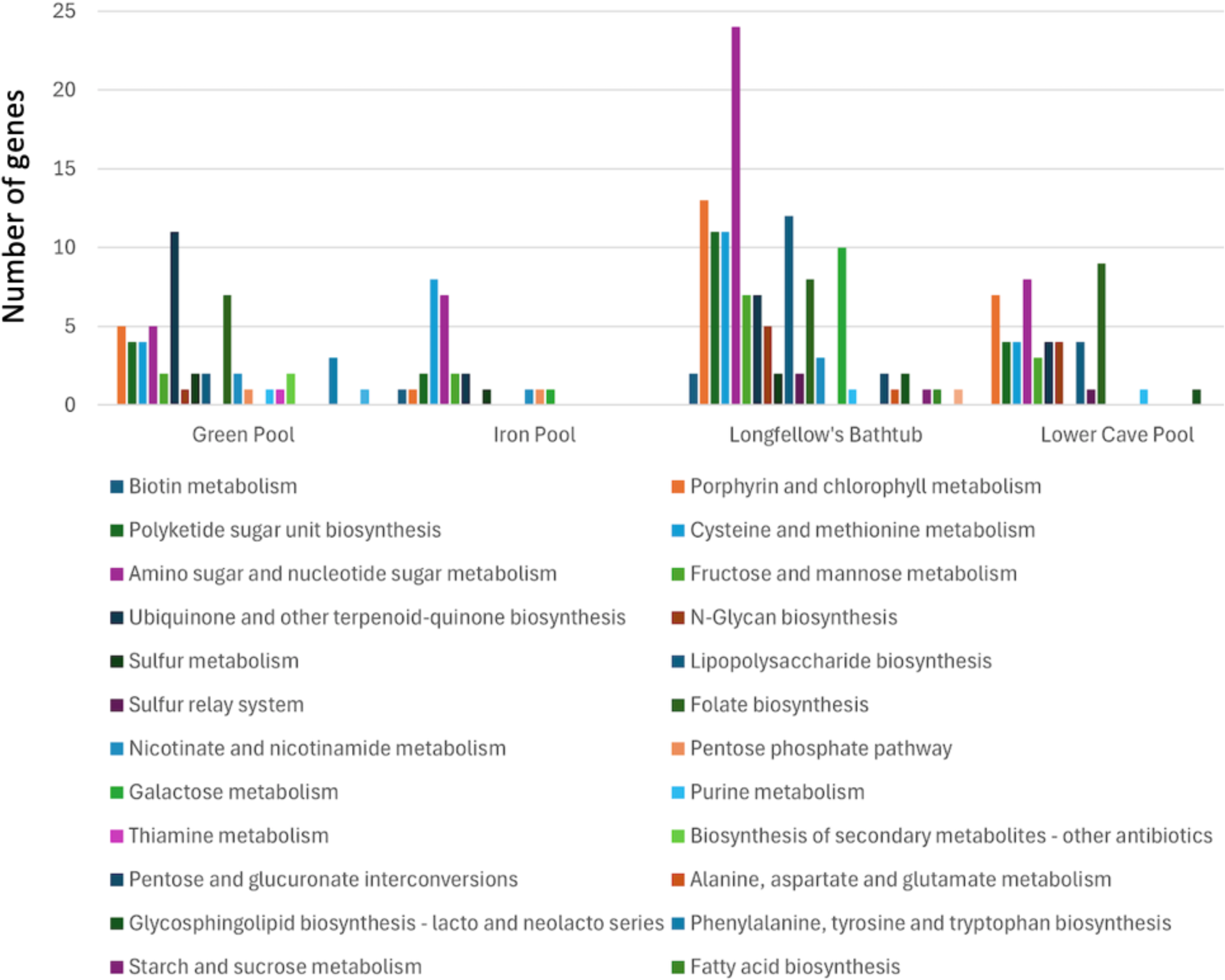
Auxiliary metabolic genes categorized by specific metabolic pathways for viral contigs of each of the four pools determined using VIBRANT.

Annotated viral MAGs of the most common virus in each pool for which a circularized genome was generated are shown in Fig. 7. The most abundant virus in Iron Pool (contig 228, Fig. 3) is an unknown phage with a genome that encodes many common viral proteins, including large terminase subunits, portal proteins, and tail length tape measure proteins. It also has large regions encoding many unknown proteins, such as the areas around 0–5 kb and 32–37 kb (Fig. 7). The most abundant virus in Lower Cave Pool is another unknown phage (contig 252) with a genome that not only shares many common phage genes with contig 228, but also encodes glycosyltransferases that are implicated in avoiding host defense mechanisms or conferring virulence factors to their host [57]. Additionally, this contig also contains two AMGs. The first encodes a D-beta-D-heptose 7-phosphate kinase / D-beta-D-heptose 1-phosphate adenosyltransferase that is likely involved in lipopolysaccharide biosynthesis, specifically the assembly of the inner core of the lipopolysaccharide inner core of gram-negative bacteria [58]. The other AMG encodes an anaerobic magnesium-protoporphyrin IX monomethyl ester cyclase which is a part of porphyrin metabolism and is utilized by anaerobes in the biosynthesis of chlorophyllide (https://www.genome.jp/entry/1.21.98.3). For the remaining two pools, the most abundant phage contigs are short and may represent fragmented genomes, so we annotated the most abundant complete circular contigs as viral MAGs. For Longfellow’s Bathtub, contig 41 is the most abundant circular contig. While it contains many recognizable phage genes, the tail and head genes are not compartmentalized like some other viruses and are grouped within each other [59–61]. There also appears to be a wide variety of genes encoding DNA/RNA and nucleotide metabolism and other functions that account for nearly half of the genes in the contig. There are also four moron, auxiliary metabolic, and host takeover genes, all encompassing genes for various transferases. Contig 138, the most abundant complete circular viral MAG in Green Lake, was assigned to the Zobellviridae family. Of note is its region around 40 Kb with significantly positive GC content and few predicted open reading frames. These regions with no predicted protein-coding regions cause a coding capacity of 88.7%, which is slightly lower than the average coding capacity of 90.45% found in the INPHARED database at the time of publication [39, 62]. Both contig 41 and contig 138 encode a glutamine-fructose-6-phosphate transaminase (isomerizing) enzyme that was predicted to be an AMG by VIBRANT.

**Fig 7.**
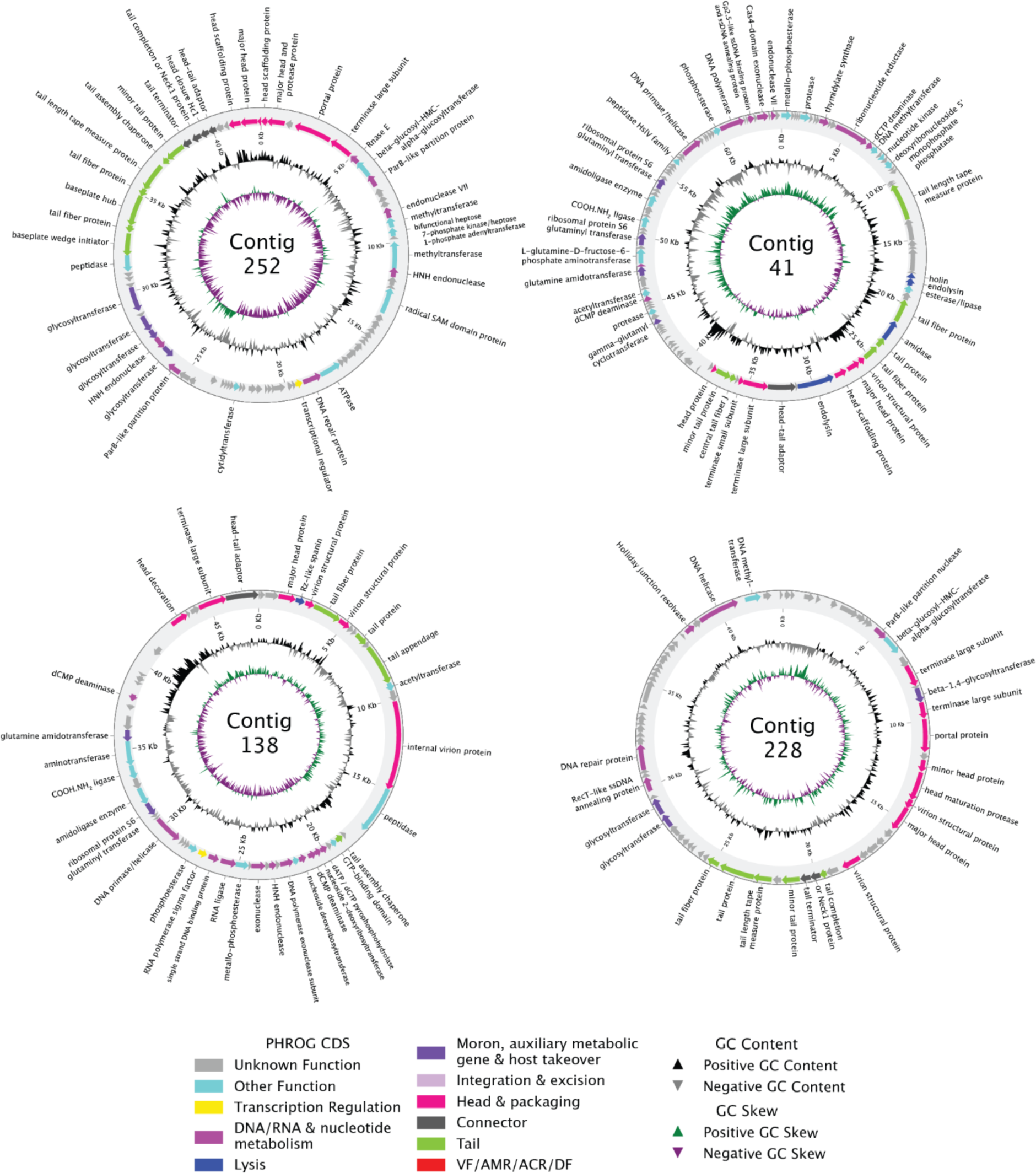
Phold-annotated viral metagenome-assembled genomes based on the high quality draft and completely circular contigs as assigned by VIBRANT of the most abundant virus from each of the four pools for which a circularized genome was obtained: Lower Cave Pool, Contig 252; Longfellow’s Bathtub, Contig 41; Green Lake, Contig 138; Iron Pool, Contig 228.

Agglomerative hierarchical clustering (Figs. 4 and 8A) showed that the viral communities from the two illuminated pools along the main tourist trail, Green Lake and Longfellow’s Bathtub, were most similar, and distinct from communities in Lower Cave Pool and Iron Pool. The Iron Pool viral community was most dissimilar from the others. However, two-way cluster analysis shows that only five abundant viral contigs are abundant in both Green lake and Longfellow’s Bathtub, and the viral communities even in the most similar sites contain many distinct viruses.

**Fig 8.**
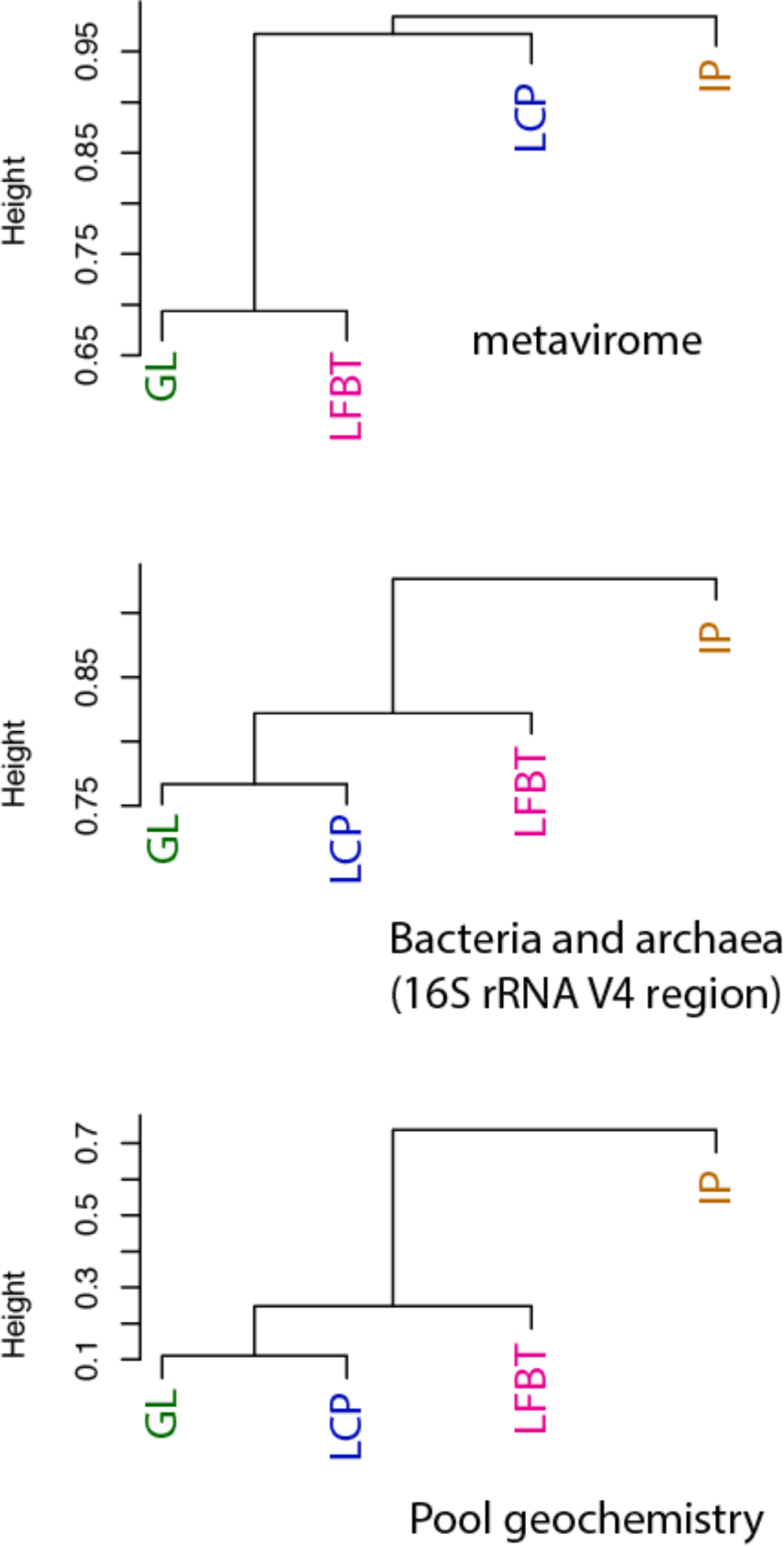
Agglomerative hierarchical clustering of A. viral communities, B. prokaryotic communities, and C. pool geochemistry among cavern pools. GL = Green Lake, LFBT = Longfellow’s Bathtub, LCP = Lower Cave Pool, IP = Iron Pool.

### Microbial Communities and Predicted Hosts

Amplicon sequencing of small subunit rRNA genes revealed a variety of bacteria and archaea in the four pools (Fig. 9). Libraries prepared using 35 PCR cycles had between 21,616 and 40,993 sequences after quality filtering and trimming, while libraries with fewer PCR cycles as well as those submitted for ‘full service’ sequencing either failed or had fewer sequences (466–19604 sequences for V4 libraries; libraries attempted with primers for archaeal and eukaryal communities failed), likely because of low template. We therefore only compared libraries generated using 35 PCR cycles prior to barcoding and the V4 primers [43], using a protocol that has been shown to produce more consistent libraries if DNA concentrations are low [47].

**Fig. 9.**
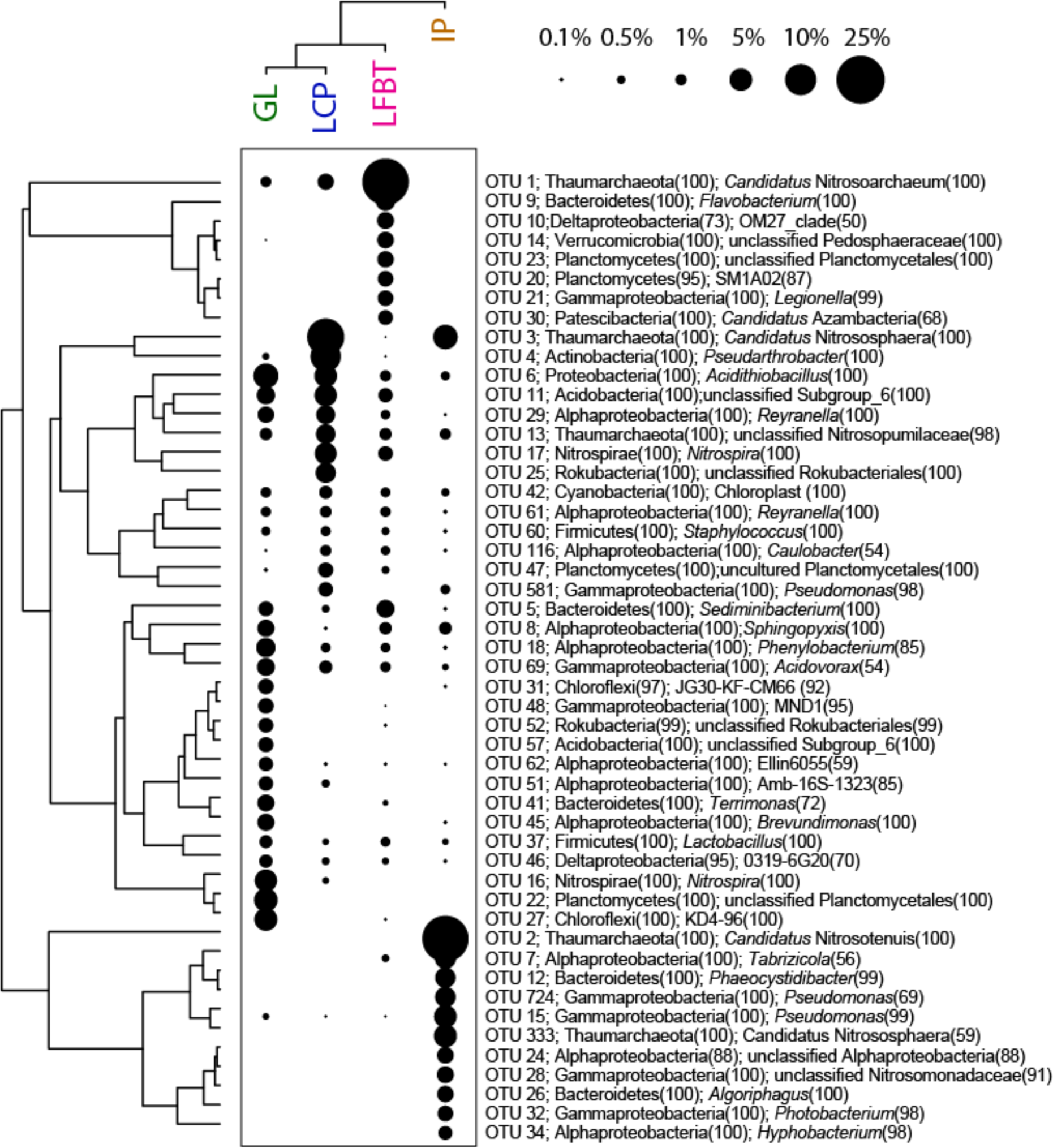
Two-way cluster analysis of microbial communities based on rRNA gene libraries. The sizes of the points scale with the relative abundance of the OTUs. Taxonomic classification are provided at the phylum and genus level, or the highest available classification for unclassified OTUs.

Based on rRNA gene libraries, the most abundant OTUs represent archaea classified as Thaumarchaeota, which comprised 20-26% of all reads and included *Candidatus* Nitrosoarchaeum, *Candidatus* Nitrosotenuis, and *Candidatus* Nitrosphaera (Fig. 9). The most common bacteria included genera *Pseudoarthrobacter*, *Nitrospira*, *Pseudomonas*, and uncultivated members of the Planctomycetales, Acidobacteria, and Chloroflexi (Fig. 7). Sequences related to cyanobacteria and chloroplasts comprised 1% or less of the reads but occurred in all four pools; BLAST searches showed the chloroplast rRNA gene sequences to be most closely related to those of higher plants. Diversity indices were similar among the four pools: Shannon indices: 3.35–4.19, inverse Simpson’s indices (1/D): 14.0–18.8; Chao 1 indices: 139–252 (Supplemental Table S2). Green Lake had the highest Shannon and inverse Simpson’s indices, while Longfellow’s Bathtub had the highest Chao1 species richness estimate.

Cluster analysis showed that the prokaryotic microbial communities of Green Lake and Lower Cave Pool were most similar to each other (Figs. 8 and 9B), and that the clustering of the bacterial communities followed the same pattern as clustering based on pool geochemical data (Fig. 8C). Iron Pool was the outlier in cluster analyses of viral and prokaryotic communities, as well as geochemistry.

Of the 1505 viral contigs generated, iPHoP was able to confidently predict hosts for 36 (Supplementary Table S3). Three were predicted to infect archaea while the rest were predicted to infect bacteria representing 30 unique host genera. Of the 30 predicted host genera, six were represented in the 16S rRNA amplicon sequences for the pools. *Prevotella*, *Nocardia*, *Mycobacterium*, *Polaromonas*, *Corynebacterium*, and *Pseudomonas* were all present in the 16S rRNA gene sequence data for the various pools, but not always corresponding to the pools where their associated viral contigs were present. For example, *Prevotella* represents one OTU of the dataset and is only present from Lower Cave Pool, but the associated viral contig has sequencing coverage from Green Lake only. Another example, *Nocardia*, is not very abundant, representing 0.003–0.007% of the pool datasets; two of the pools it was present in correspond to the two pools for which the viral contig had coverage. Based on the relative abundance of these genera in the rRNA gene libraries, potential viral hosts include abundant and rare members of each pool community (Fig. 10, Supplemental Table S3). Likewise, OTUs in the same families as these predicted hosts also occur among the abundant and rare members of the communities. Because many of the microbial OTUs in the dataset are from unnamed groups that are often only know from rRNA gene phylogenies, these cannot easily be linked to potential hosts either because they do not have representatives with genome or metagenome-assembled genomes, or because predicted hosts classified with the genome taxonomy database (GTDB) system used by iPHoP cannot often be linked to rRNA gene classification using the Silva taxonomy.

**Fig 10.**
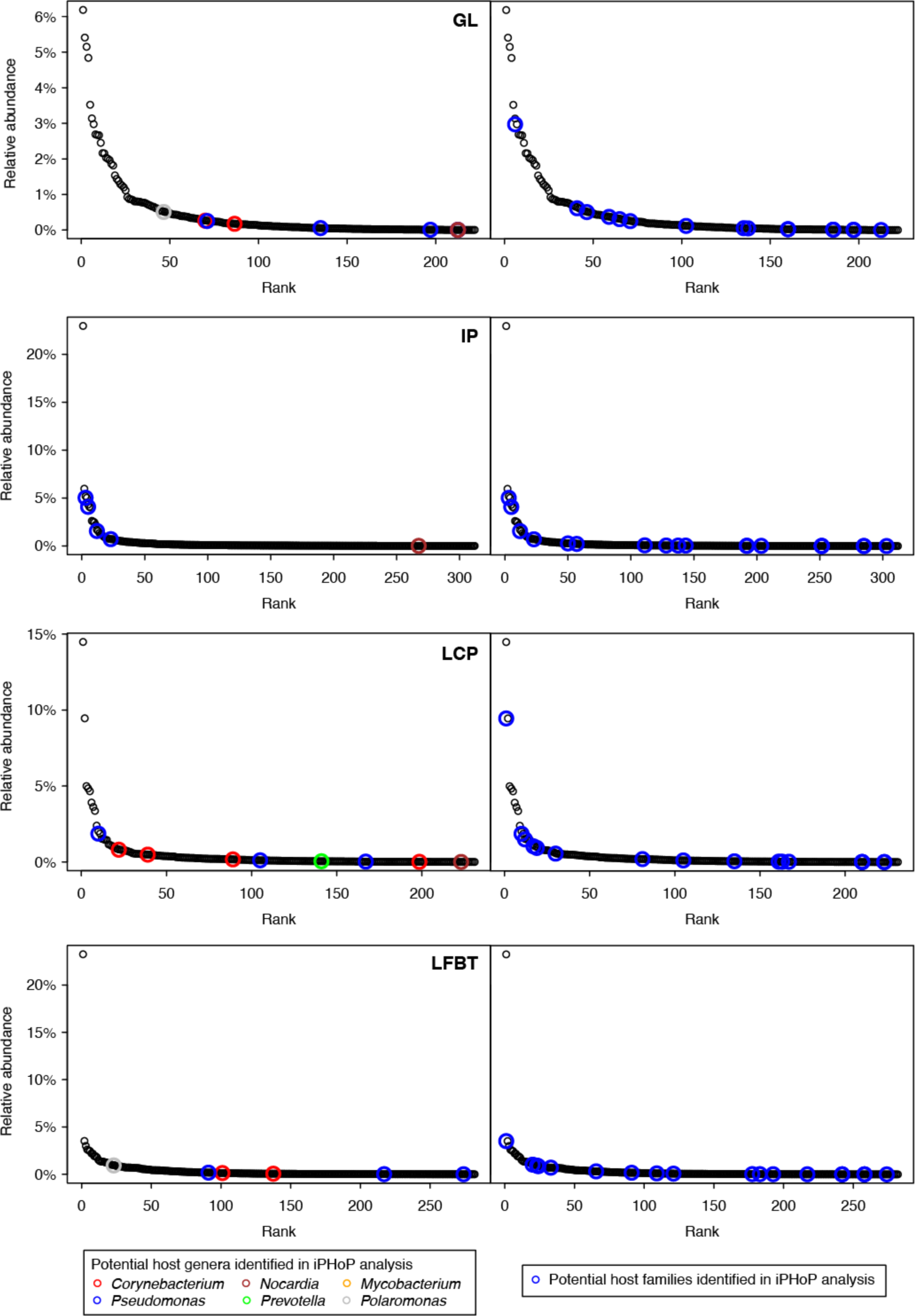
Rank abundance curves of rRNA gene amplicon libraries, with circles indicating possible hosts as determined using iPHoP. The left panels show the rank of OTUs classified as genera identified as potential hosts by iPHoP, and the right panels show the rank of OTUs classified as families identified as potential hosts.

## Discussion

The first hypothesis tested in this study was supported by the data. Microscopic counts of bacteria averaged ∼1 × 10^4^ cells mL^-1^ and viral counts averaged ∼2 × 10^5^ VLPs mL^-1^. These data are in rough agreement with those from marine and other environments, and thus support the hypothesis that virus abundance would be an order of magnitude higher than microbial abundance. The second hypothesis was that cave pools would contain novel phages. Of the 1505 high-confidence viral contigs, most of them represent phages, but only 418 clustered with known viruses. Therefore, in support of the project’s second hypothesis, most (80%, Fig. 5) of the viruses in Carlsbad Cavern pools are therefore novel and add to our knowledge of “viral dark matter”. The third hypothesis, that viral communities of pools in developed cave areas would differ from those in undeveloped cave areas, was also supported. Green Lake and Longfellow’s Bathtub share characteristics of heavy tourist traffic and artificial lighting (and associated lampenflora microbes including photosynthetic Cyanobacteria). Somewhat surprisingly, while the viral communities clustered as hypothesized, the prokaryotic communities clustered differently and instead appear to reflect the water chemistry of the pools (Fig. 8).

Some of the high confidence viral contigs were classified to the family level utilizing graphanalyzer in conjunction with vConTACT2 and the INPHARED database. Those that could be classified to the family level belong to the following viral families: Autographiviridae (N=3), Zobellviridae (N=6), Podoviridae (N=1), Mesyanzhinoviridae (N=3), Suoliviridae (N=1), and Casjensviridae (N=1) all of which belong to the class Caudoviricetes. An additional 53 viral contigs were also classified to the Caudoviricetes class but did cluster tightly enough to inherit family classification. The Autographiviridae are characterized by podovirus morphology and T7-like RNA polymerases [63].

The Siphoviridae, Podoviridae, Myoviridae, Herelleviridae, Demerecviridae, and Autographyiridae are all members of the morphologically-based viral order Caudovirales, all of which are phages with tails of various lengths, and these comprised 94% of the sequences in this study that have homology to known viruses, although a very small proportion of total viral contigs. The Caudovirales are evidently a highly successful viral group, comprising ∼90% of sequences among ∼1200 viral sequences linked to host cells genomes derived from a variety of environments [64]. Members of the Caudovirales infect many bacterial phyla, including Cyanobacteria, as well as archaea [65, 66]. It is thus not surprising to find a diversity of Caudovirales in the cave pools, especially the lighted ones where Cyanobacteria may occur.

The Phycodnaviridae are known to infect eukaryotic algae [67], which is consistent with the artificial illumination and green biofilms in Green Lake and Longfellow’s Bathtub, as lampenflora in other lighted parts of the cave have abundant Chlorophyta and Ochrophyta [5]. Also, at least some of the members of this family are giant viruses [68]. Some of the Mimiviridae are giant viruses that infect amoebas and that can be up to 500 nm in diameter [69]. However, further study is needed to determine whether the sequences affiliated to these two viral families are actually giant viruses. If so, then they apparently passed through the 0.2-µm filter at the time of sampling.

While the majority of phages and archaeal viruses are thought to be DNA viruses [70], RNA phages are also known [25], but any of these that exist in the cave pools would be missed by the methods used here. Of the viral sequences detected in this study, the majority are unrelated to previously sequenced viruses and furthermore, a large proportion of the putative genes (ORFs) in this study have no homology to database genes encoding known function—that is, they are ORFans (open reading frames lacking homologues in existing databases). Therefore, this study primarily indicates the presence of diverse, mostly previously unknown viruses, but it provides only glimpses of their roles in the environment. Another limitation is that the viromes of these four pools are likely not representative of all cave pools or all cave environments. The study has merely scratched the subsurface; it is hoped that further surveys of cave viruses will follow.

This first ever metagenomic study of cave viruses revealed diverse and novel viruses that infect cave pool bacteria. The discovery of these hitherto unstudied biological entities in a cave should pave the way for future studies of viruses in cave and karst environments. Metagenomic surveys of viruses should be extended to cave sediments and biofilms in addition to caves with varied physical and chemical conditions as these are apt to yield more new viruses. Studies that link viruses to their specific hosts and that determine the possible negative and positive effects on those hosts will take cave virus studies to a new level. Better understanding of the role of viruses in caves may even have implications for management of cave resources. For example, efforts could be made to cultivate viruses that infect algae and cyanobacteria for the purpose of controlling lampenflora populations.

## Supplementary Information

## Acknowledgements

We thank the U.S. National Park Service’s Carlsbad Caverns National Park personnel for access to Carlsbad Cavern. We thank Ellen Trautner for help in site selection, Kasandra Velarde for help in sampling, and Alex Probst for insightful advice and discussions about metavirome preparation and analysis. Samples were collected under research permit number CAVE-2020-SCI-0002.

## Author Contributions

TLK conceived the study and obtained funding. JU collected samples and performed laboratory steps of the study. JU, NEJ, and DSJ carried out bioinformatic analyses. All authors contributed to writing the manuscript and approved the final version.

## Data Availability

Data are provided within the manuscript or supplementary information. Raw sequence data are available under BioProject accession number PRJNA1159728 (https://www.ncbi.nlm.nih.gov/bioproject/). High confidence viral contigs from the metavirome co-assembly are available at https://doi.org/10.6084/m9.figshare.26993332.

## References

1. Palmer AN (2007) Cave Geology. Dayton, Ohio: Cave Books, Dayton, Ohio.

2. Barton, HA. (2006) Introduction to cave microbiology: a review for the non-specialist. J. Cave Karst Studies 68:43–54.

3. Barton HA, Northup DE (2007) Geomicrobiology in cave environments: past, current and future perspectives. Journal of Cave and Karst Studies 69:163–178

4. Snider JR, Goin C, Miller RV, Boston PJ, Northup DE (2009) Ultraviolet radiation sensitivity in cave bacteria: evidence of adaptation to the subsurface? Internatl J Speleol 38:11–22.

5. Havlena Z, Kieft TL, Veni G, Horrocks RD, Jones DS, Liu S-J (2021) Lighting effects on the development and diversity of photosynthetic biofilm communities in Carlsbad Cavern, New Mexico. Appl Environ Microbiol 87:e02695–20. 10.1128/AEM.02695-20

6. Read, KJH, Melim LA, Winter AS, Northup DE (2021) Bacterial diversity in vadose zone cave pools: evidence for isolated ecosystems. J Cave Karst Studies 83:163–188. 10.4311/2020MB0120

7. Constantine DG, Woodall DF (1964) Latent infection of Rio Bravo virus in salivary glands of bats. Public Health Reports 79:1033–1039.

8. Constantine DG, Tierkel ES, Kleckner MD, Hawkins DM (1968) Rabies in New Mexico cavern bats. Public Health Reports 83:303–316

9. Ghaly TM, Focardi A, Elbourne LDH, Sutcliffe B, Humphreys WF, Jaschke PR, Tetu SG, Paulsen IT (2024) Exploring virus-host-environment interactions in a chemotrophic-based underground estuary. Environ Microbiome 19:9. 10.1186/s40793-024-00549-6

10. Dávila-Ramos S, Castelán-Sánchez HG, Martínez-Ávila L, Sánchez-Carbente MDR, Peralta R, Hernández-Mendoza A, Dobson ADW, Gonzalez RA, Pastor N, Batista-García RA. 2019. A review on viral metagenomics in extreme environments. Front Microbiol 10:2403. 10.3389/fmicb.2019.02403

11. Gil JF, Mesa V, Estrada-Ortiz N, Lopez-Obando M, Gómez A, Plácido J (2021) Viruses in extreme environments, current overview, and biotechnological potential. Viruses-Basel 13:81. 10.3390/v13010081

12. Weinbauer MG (2004) Ecology of prokaryotic viruses. FEMS Microbiol Rev 28:127–181. 10.1016/j.femsre.2003.08.001

13. Comeau AM, Hatfull GF, Krisch HM, Lindell D, Mann NH, Prangishvili D (2008) Exploring the prokaryotic virosphere. Res Microbiol 159:306–13 10.1016/j.resmic.2008.05.001

14. Wilhelm SW, Suttle CA (1999) Viruses and nutrient cycles in the sea: viruses play critical roles in the structure and function of aquatic food webs. BioScience 49:781–788. 10.2307/1313569

15. Suttle CA (2007) Marine viruses–major players in the global ecosystem. Nat Rev Microbiol 5:801–12. 10.1038/nrmicro1750

16. Rohwer F, Prangishvili D, Lindell D (2009) Roles of viruses in the environment. Environmental Microbiology 11:2771–2774. 10.1111/j.1462-2920.2009.02101.x

17. Sullivan MB, Krastins B, Hughes JL, Kelly L, Chase M, Sarracino D, Chisholm SW (2009) The genome and structural proteome of an ocean siphovirus: a new window into the cyanobacterial ‘mobilome’. Environ Microbiol 11:2935–2951. 10.1111/j.1462-2920.2009.02081.x

18. Allen MJ, Forster T, Schroeder DC, Hall M, Roy D, Ghazal P, Wilson WH (2006) Locus-specific gene expression pattern suggests a unique propagation strategy for a giant algal virus. J Virol 80:7699–7705. 10.1128/JVI.00491-06

19. Hayes S, Mahony J, Nauta A, van Sinderen D (2017) Metagenomic approaches to assess bacteriophages in various environmental niches. Viruses-Basel 9:127. 10.3390/v9060127

20. Lloyd KG, Steen AD, Ladau J, Yin YQ, Crosby L (2018) Phylogenetically novel uncultured microbial cells dominate Earth microbiomes. MSystems 3:e00055–18. 10.1128/mSystems.00055-18

21. Steen AD, Crits-Christoph A, Carini P, DeAngelis KM, Fierer N, Lloyd KG, Thrash JC (2019) High proportions of bacteria and archaea across most biomes remain uncultured. ISME J 13:3126–3130, 10.1038/s41396-019-0484-y

22. Rappé MS, Giovannoni SJ (2003) The uncultured microbial majority. Ann Rev Microbiol 57:369–394. 10.1146/annurev.micro.57.030502.090759

23. Waldron D (2015) Sorting out viral dark matter. Nature Rev Microbiol 13:526–527. 10.1038/nrmicro3541

24. Breitbart M, Salamon P, Andresen B, Mahaffey JM, Segall AM, Mead D, Azam F, Rohwer F (2002) Genome analysis of uncultured marine viral communities. Proc Natl Acad Sci 99:14250–14255. 10.1073/pnas.202488399

25. Callanan J, Stockdale SR, Shkoporov A, Draper LA, Ross, RP, Hill C. (2018) RNA phage biology in a metagenomic era. Viruses-Basel 10:386. 10.3390/v10070386

26. John SG, Mendez CB, Deng L, Poulos B, Kauffman AKM, Kern S, Brum J, Polz MF, Boyle EA, Sullivan MB (2011) A simple and efficient method for concentration of ocean viruses by chemical flocculation. Environ Microbiol Reports 3:195–202. 10.1111/j.1758-2229.2010.00208.x

27. Hobbie JE, Daley RJ, Jasper S (1977) A method for counting bacteria on Nucleporefilters. Appl Environ Microbial 33: 1225–1228.

28. Noble R, Fuhrman J (1998) Use of SYBR Green I for rapid epifluorescence counts of marine viruses and bacteria. Aquat Microb Ecol 14:113–118. 10.3354/ame014113

29. Shibata A, Yoichi G, Saito H, Kikuchi T, Toda T, Taguchi S (2006) Comparison of SYBR Green I and SYBR Gold stains for enumerating bacteria and viruses by epifluorescence microscopy. Aquat Microb Ecol 43:223–231. 10.3354/ame043223

30. Hurwitz BL, Deng L, Poulos BT, Sullivan MB. (2013) Evaluation of methods to concentrate and purify ocean virus communities through comparative, replicated metagenomics. Environ Microbiol 15:1428–1440. 10.1111/j.1462-2920.2012.02836.x

31. Joshi N, Fass J. 2011. Sickle: A sliding-window, adaptive, quality-based trimming tool for FastQ files vVersion 1.33. https://github.com/najoshi/sickle

32. Martin M (2011) Cutadapt removes adapter sequences from high-throughput sequencing reads. EMBnet.journal 17:10–12. 10.14806/ej.17.1.200

33. Nurk S, Meleshko D, Korobeynikov A, Pevzner PA (2017) metaSPAdes: a new versatile metagenomic assembler. Genome Res 27:824–834. 10.1101/gr.213959.116

34. Guo JR, Bolduc B, Zayed AA, Varsani A, Dominguez-Huerta G, Delmont TO, Pratama AA, Gazitúa MC, Vik D, Sullivan MB, Roux S (2021) VirSorter2: a multi-classifier, expert-guided approach to detect diverse DNA and RNA viruses. Microbiome 9:37. 10.1186/s40168-020-00990-y

35. Ren J, Ahlgren NA, Lu YY, Fuhrman JA, Sun F (2017) VirFinder: a novel k-mer based tool for identifying viral sequences from assembled metagenomic data. Microbiome 5:69. 10.1186/s40168-017-0283-5

36. Jang HB, Bolduc B, Zablocki O, Kuhn JH, Roux S, Adriaenssens EM, Brister JR, Kropinski AM, Krupovic M, Lavigne R, Turner D, Sullivan MB (2019) Taxonomic assignment of uncultivated prokaryotic virus genomes is enabled by gene-sharing networks. Nat Biotechnol 37:632–639. 10.1038/s41587-019-0100-8

37. Hyatt D, Chen G-L, LoCascio PF, Land ML, Larimer FW, Hauser LJ. (2010) Prodigal: prokaryotic gene recognition and translation initiation site identification. BMC Bioinformatics 11:119. 10.1186/1471-2105-11-119

38. Shannon P, Markiel A, Ozier O, Baliga NS, Wang JT, Ramage D, Amin N, Schwikowski B, Ideker T (2003) Cytoscape: a software environment for integrated models of biomolecular interaction networks. Genome Res 13:2498–2504. 10.1101/gr.1239303

39. Cook R, Brown N, Redgwell T, Rihtman T, Barnes M, Clokie M, Stekel DV, Hobman J, Jones MA, Millard A (2021) INfrastructure for a PHage REfrerence Database: identification of large-scalebiases in the current collection of cultured phage genomes. Phage 2:214–223. 10.1089/phage.2021.0007

40. Pandolfo M, Telatin A, Lazzari G, Adriaenssens EM, Vitulo N (2022) MetaPhage: an Automated Pipeline for Analyzing, Annotating, and Classifying Bacteriophages in Metagenomics Sequencing Data. mSystems 7:e00741–22. 10.1128/msystems.00741-22

41. Roux S, Camargo AP, Coutinho FH, Dabdoub SM, Dutihl BE, Nayfach S, Tritt A (2023) iPHoP: An integrated machine learning framework to maximize host prediction for metagenome-derived viruses of archaea and bacteria. PLOS Biol 21:e3002083. 10.1371/journal.pbio.3002083

42. Kieft K, Zhou Z, Anantharaman (2020) VIBRANT: an automated recovery, annotation and curation of microbial viruses, and evaluation of viral community function from genomic sequences. Microbiome 8:90. 10.1186/s40168-020-00867-0

43. Walters W, Hyde ER, Berg-Lyons D, Ackerman G, Humphrey G, Parada A, Gilbert JA, Jannson JK, Caporaso JG, Fuhrman JA, Apprill A, Knight R. (2016) Improved bacterial 16S rRNA (V4 and V4-5) and fungal internal transcribed spacer marker gene primers used for microbial community surveys. mSystems 1:e00009–15.

44. Stahl DA, Amann RI (1991) Development and application of nucleic acid probes in bacterial systematics, p 205–248. In Stackebrandt E, Goodfellow M (ed), Nucleic acid techniques in bacterial systematics. John Wiley & Sons Ltd, Chichester, UK.

45. Caporaso JG, Lauber CL, Walters WA, Knight R (2011) Global patterns of 16S rRNA diversity at a depth of millions of sequences per sample. Proc Natl Acad Sci 108(supplement 1):4516–4522. 10.1073/pnas.1000080107

46. Gohl DM, MacLean A, Hauge A, Becker A, Walek D, Beckman KB (2016) An optimized protocol for high-throughput amplicon-based microbiome profiling. 10.1038/protex.2016.030

47. Jones DS, Lapenko, KA, Wenz ZJ, Olson MC, Roepke EW, Sadowsky MJ, Novak PJ, Bailey JV (2017) Novel microbial assemblages dominate weathered sulfide-bearing rock from copper-nickel deposits in the Duluth complex, Minnesota, USA. Appl Environ Microbiol 83:e00909–17. 10.1128/AEM.00909-17

48. Zhang J, Kobert K, Flouri T, Stamatakis A (2014) PEAR: a fast and accurate Illumina Paired-End reAd mergeR. Bioinformatics30:614–620. 10.1093/bioinformatics/btt593

49. Edgar RC (2013) Uparse: highly accurate OTU sequences from microbial amplicon reads. Nature Methods 10:996–998. 10.1038/NMETH.2604

50. Pruesse E, Quast C, Knittel K, Fuchs BM, Ludwig WG, Peplies J, Gloeckner FO (2007) SILVA: a comprehensive online resource for quality checked and aligned ribosomal RNA sequence data compatible with ARB Nucleic Acids Res 35:7188–7196. 10.1093/nar/gkm864

51. Schloss PD, Westcott SL, Ryabin T, Hall JR, Hartmann M, Hollister EB, Lesniewski RA, Oakley BB, Parks DH, Robinson CJ, Sahl JW, Stres B, Thallinger GG, Van Horn DJ, Weber CF (2009) Introducing mothur: Open-source, platform-independent, community-supported software for describing and comparing microbial communities. Appl Environ Microbiol 75:7537–7541. 10.1128/AEM.01541-09

52. R Core Team (2023) R: A Language and Environment for Statistical Computing. R Foundation for Statistical Computing, Vienna. https://www.R-project.org/

53. Oksanen FJ, et al. (2017) Vegan: Community Ecology Package. R package Version 2.4–3. https://CRAN.R-project.org/package=vegan

54. Shannon CE (1948) A mathematical theory of communication. Bell System Technical J 27:379–423. 10.1002/j.1538-7305.1948.tb01338.x

55. Hunter, PR, Gaston, MA (1988). Numerical index of the discriminatory ability of typing systems: an application of Simpson’s index of diversity. J Clin Microbiol 26: 2465– 2466. 10.1128/JCM.26.11.2465-2466

56. Chao A., Chiu C. (2016) Species Richness: Estimation and Comparison. In: Wiley StatsRef: Statistics Reference Online. pp 1–26 10.1002/9781118445112.stat03432.pub2

57. Markine-Goriaynoff N, Gillet L, Van Etten JL, Korres H, Verma N, Vanderplasschen A (2004) Glyosyltransferases encoded by viruses. J Gen Virol 85:2741–2754. 10.1099/vir.0.80320-0

58. Wang L, Huang H, Nguyen HH, Allen KN, Mariano PS, Dunaway-Mariano D (2010) Divergence of biochemical function in the HAD superfamily: D-glycero-D-manno-Heptose-1,7-bisphosphate phosphatase (GmhB) Biochem 49:1072–1081. 10.1021/bi902018y

59. Pedulla ML, Ford ME, Houtz JM, Karthikeyan T, Wadsworth C, Lewis JA, Jacobs-Sera D, Falbo J, Gross J, Pannunzio NR, Brucker W, Kumar V, Kandasamy J, Keenan L, Bardarov S, Kriakov J, Lawrence JG, Jacobs WR, Hendrix RW, Hatfull GF (2003) Origins of highly mosaic mycobacteriophage genomes. Cell 113: 171–182. 10.1016/S0092-8674(03)00233-2

60. Xu J, Hendrix RW, Duda RL (2004) Conserved translational frameshift in dsDNA bacteriophage tail assembly genes. Molec Cell 16:11–21. 10.1016/j.molcel.2004.09.006

61. Hatful GF (2008) Bacteriophage genetics. Curr Opin Microbiol 11:447–453. 10.1016/j.mib.2008.09.004

62. Turner D, Adriaenssens EM, Tolstoy I, Kropinski AM (2021) Phage annotation guide: guidelines for assembly and high-quality annotation. Phage 2:170–182. 10.1089/phage.2021.001

63. Boeckman J, Korn A, Yao G, Ravindran A, Gonzalez C, Gill J (2022) Sheep in wolves’ clothing: temperate T7-like bacteriophages and the origins of the *Autographiviridae*. Virol 568:86–100. 10.1016/j.virol.2022.01.013

64. Roux S, Hallam SJ, Woyka T, Sullivan MB (2015) Viral dark matter and virus-host interactions resolved from publicly available microbial genomes. eLife 4:e08490. 10.7554/eLife.08490

65. Ackermann HW (1998) Tailed bacteriophages: the order *Caudovirales*. Adv Virus Res 51:135–201.

66. Maniloff J, Ackerman HW (1998) Taxonomy of bacterial viruses: establishment of tailed virus genera and the order *Caudovirales* Arch Virol 143:2051–2063. 10.1007/s007050050442

67. Van Etten JL, Graves MV, Müller DG, Boland W, Delaroque N. (2002) Phycodnaviridae – large DNA algal viruses. Arch Virol 147:1479–1516. 10.1007/s00705-002-0822-6

68. Schulz F, Abergel C, Woyke T (2022) Giant virus biology and diversity in the era of genome-resolved metagenomics. Nat Rev Microbiol 20:721–736. 10.1038/s41579-022-00754-5

69. Colson P, La Scola B, Levasseur A, Caetano-Anoliés G, Raoult D (2017) Mimivirus: leading the way in the discovery of gian viruses of amoebae. Nat Rev Microbiol 15:243–254. https://10.1038/nrmicro.2016.197

70. Ackermann HW (2007) 5500 phages examined in the electron microscope. Arch Virol 152(2):227–243 10.1007/s00705-006-0849-1

