## Supplementary Information for "Cave pools in Carlsbad Caverns National Park contain diverse bacteriophage communities and novel viral sequences"

**Supplemental Table S1** Elemental analysis of pool water by ICP-MS.

| Element (mg $L^{-1}$) | Longfellow's Bathtub | Green Lake | Lower Cave Pool | Iron Pool |
|---|---|---|---|---|
| Aluminum | BD[a] | 0.0007 | BD[a] | BD[a] |
| Antimony | BD[a] | BD[a] | BD[a] | BD[a] |
| Arsenic | BD[a] | 0.0010 | 0.0011 | BD |
| Barium | 0.089 | 0.259 | 0.311 | 0.090 |
| Beryllium | BD[a] | BD[a] | BD[a] | BD[a] |
| Boron | 0.073 | 0.061 | 0.047 | 0.492 |
| Cadmium | BD[a] | BD[a] | BD[a] | BD[a] |
| Calcium | ND[b] | ND[b] | 19.5 | 31.2 |
| Chromium | BD[a] | BD[a] | 0.0010 | BD[a] |
| Cobalt | BD[a] | BD[a] | BD[a] | BD[a] |
| Copper | BD[a] | 0.0026 | BD[a] | BD[a] |
| Iron | BD[a] | BD[a] | BD[a] | BD[a] |
| Lead | BD[a] | BD[a] | BD[a] | BD[a] |
| Lithium | 0.005 | 0.005 | 0.005 | 0.394 |
| Magnesium | BD[a] | BD[a] | 36.6 | 1850 |
| Manganese | BD[a] | BD[a] | BD[a] | BD[a] |
| Molybdenum | BD[a] | 0.001 | 0.001 | 0.193 |
| Nickel | BD[a] | 0.0012 | BD[a] | BD[a] |
| Potassium | ND[b] | ND[b] | 0.53 | 372 |
| Selenium | BD[a] | 0.001 | 0.001 | 0.194 |
| Silicon | 18.9 | 8.99 | 9.29 | 1.21 |
| Silver | BD[a] | BD[a] | BD[a] | BD[a] |
| Strontium | 0.227 | 0.115 | 0.260 | 0.148 |
| Thallium | BD[a] | BD[a] | BD[a] | BD[a] |
| Thorium | BD[a] | BD[a] | BD[a] | BD[a] |

| | | | | |
|---|---|---|---|---|
| Tin | BD[a] | 0.0012 | 0.008 | BD[a] |
| Titanium | BD[a] | 0.001 | 0.001 | BD[a] |
| Uranium | BD[a] | 0.0013 | 0.0015 | BD[a] |
| Vanadium | 0.0142 | 0.0096 | 0.0044 | BD[a] |
| Zinc | 0.0213 | 0.0076 | 0.0128 | BD[a] |

[a]BD, below detection limit
[b]ND, no data

**Supplemental Table S2 Diversity indices for prokaryotes based on amplicon sequencing data.**

| Index | Lower Cave Pool | Longfellow's Bathtub | Iron Pool | Green Lake |
|---|---|---|---|---|
| Shannon | 3.38 | 4.03 | 3.67 | 4.19 |
| Inverse Simpson's (1/D) | 17.1 | 17.3 | 14.0 | 18.8 |
| Chao1 | 139 | 252 | 235 | 187 |

**Supplemental Table S3** List of potential cellular host species based on iPHoP analysis.

| Contig | Length (bp) | Host classification: domain (d), phylum (p), class (c), order (o), family (f), genus (g) | Confi-dence score | Method(s), with scores from individual method(s) |
|---|---|---|---|---|
| 86 | 54743 | d: Bacteria, d: Desulfobacterota, c: Desulfobaccia, o: Desulfobaccales, f: 0-14-0-80-60-11, g: 0-14-0-80-60-11 | 91.6 | BLAST, 93.90 |
| 161 | 46316 | d: Bacteria, d: Acidobacteriota, c: Thermoanaerobaculia, o: Gp7-AA8, f: Gp7-AA8, g: JADGNZ01 | 90.8 | BLAST, 93.20 |
| 167 | 46116 | d: Bacteria, d: Bacillota_A, c: Clostridia, o: Oscillospirales, f: Ruminococcaceae, g: *Ruminiclostridium*_E | 98.5 | CRISPR, 98.90 |
| 252 | 42485 | d: Bacteria, d: Bacillota_A, c: Clostridia, o: Lachnospirales, f: Lachnospiraceae, g: *Eubacterium*_G | 91.0 | iPHoP-RF, 93.40 |
| 356 | 39871 | d: Bacteria, d: Bacillota_A, c: Clostridia, o: Lachnospirales, f: Lachnospiraceae, g: *Blautia* | 90.3 | iPHoP-RF, 92.80 |
| 542 | 32336 | d: Bacteria, d: Actinomycetota, c: Acidimicrobiia, o: Acidimicrobiales, f: Ilumatobacteraceae, g: UBA2093 | 94.9 | BLAST, 96.40<br>iPHoP-RF, 62.50 |
| 608 | 29806 | d: Bacteria, d: Pseudomonadota, c: Gammaproteobacteria, o: Pseudomonadales, f: Pseudomonadaceae, g: *Pseudomonas*_E | 92.2 | iPHoP-RF, 94.40 |
| 898 | 20594 | d: Bacteria, d: Pseudomonadota, c: Gammaproteobacteria, o: Burkholderiales, f: Burkholderiaceae, g: *Paraburkholderia* | 93.5 | iPHoP-RF, 95.40 |
| 1250 | 14773 | d: Bacteria, d: Bacillota, c: Bacilli, o: Lactobacillales, f: Carnobacteriaceae, g: *Atopostipes* | 90.3 | iPHoP-RF, 92.80 |
| 1752 | 11111 | d: Bacteria, d: Actinomycetota, c: Actinomycetia, o: Mycobacteriales, f: Mycobacteriaceae, g: *Mycobacterium* | 92.6 | iPHoP-RF, 94.70 |
| 1873 | 10561 | d: Bacteria, d: Bacillota, c: Bacilli, o: Bacillales, f: DSM-18226, g: *Neobacillus* | 91.0 | iPHoP-RF, 93.40 |
| 2975 | 6897 | d: Bacteria, d: Actinomycetota, c: Actinomycetia, o: Mycobacteriales, f: Mycobacteriaceae, g: *Mycobacterium* | 93.9 | iPHoP-RF, 95.70 |
| 3293 | 6352 | d: Bacteria, d: Bacillota_A, c: Clostridia, o: Oscillospirales, f: Acutalibacteraceae, g: *Ruminococcus*_E | 98.1 | CRISPR, 98.60<br>iPHoP-RF, 53.60 |
| 4088 | 5251 | d: Bacteria, d: Actinomycetota, c: Actinomycetia, o: Propionibacteriales, f: Propionibacteriaceae, g: WQYJ01 | 90.3 | iPHoP-RF, 92.80 |
| 4177 | 5166 | d: Bacteria, d: Pseudomonadota, c: Gammaproteobacteria, o: Burkholderiales, f: Burkholderiaceae_B, g: *Polaromonas* | 93.9 | iPHoP-RF, 95.70 |
| 4555 | 4811 | d: Bacteria, d: Pseudomonadota, c: Gammaproteobacteria, o: Enterobacterales_A, f: Shewanellaceae, g: *Shewanella* | 91.8 | iPHoP-RF, 94.10 |

| | | | | |
|---|---|---|---|---|
| 5026 | 4417 | d: Bacteria, d: Bacteroidota, c: Bacteroidia, o: Flavobacteriales, f: Flavobacteriaceae, g: Winogradskyella | 90.3 | iPHoP-RF, 92.80 |
| 5066 | 4389 | d: Bacteria, d: Bacteroidota, c: Ignavibacteria, o: Ignavibacteriales, f: Melioribacteraceae, g: DSXH01 | 90.9 | BLAST, 93.30<br>iPHoP-RF, 51.50 |
| 5541 | 4044 | d: Bacteria, d: Pseudomonadota, c: Alphaproteobacteria, o: Rhizobiales, f: Xanthobacteraceae, g: *Bradyrhizobium* | 95.3 | iPHoP-RF, 96.70 |
| 5885 | 3846 | d: Bacteria, d: Actinomycetota, c: Actinomycetia, o: Mycobacteriales, f: Mycobacteriaceae, g: *Corynebacterium* | 90.3 | iPHoP-RF, 92.80 |
| 5931 | 3820 | d: Bacteria, d: Pseudomonadota, c: Gammaproteobacteria, o: Pseudomonadales, f: Moraxellaceae, g: *Psychrobacter* | 90.7 | iPHoP-RF, 93.10 |
| 6086 | 3742 | d: Bacteria, d: Chloroflexota, c: Anaerolineae, o: Aggregatilineales, f: Phototrophicaceae, g: OLB13 | 92.2 | iPHoP-RF, 94.40 |
| 6094 | 3736 | d: Bacteria, d: Patescibacteria, c: Paceibacteria, o: UBA9983_A, f: UBA1539_A, g: UBA1550 | 93.1 | iPHoP-RF, 95.10 |
| 6094 | 3736 | d: Bacteria, d: Patescibacteria, c: Paceibacteria, o: UBA9983_A, f: SBAW01, g: SBAW01 | 91.8 | iPHoP-RF, 94.10 |
| 6772 | 3447 | d: Bacteria, d: Actinomycetota, c: Actinomycetia, o: Mycobacteriales, f: Mycobacteriaceae, g: *Corynebacterium* | 95.3 | iPHoP-RF, 96.70 |
| 6997 | 3359 | d: Archaea, d: Thermoproteota, c: Bathyarchaeia, o: B26-1, f: BA1, g: BA2 | 93.5 | iPHoP-RF, 95.40 |
| 6997 | 3359 | d: Archaea, d: Thermoproteota, c: Bathyarchaeia, o: B26-1, f: BA1, g: BA1 | 92.6 | iPHoP-RF, 94.70 |
| 6997 | 3359 | d: Archaea, d: Thermoproteota, c: Bathyarchaeia, o: B26-1, f: BA1, g: Kmv02 | 90.7 | iPHoP-RF, 93.10 |
| 9110 | 2721 | d: Bacteria, d: Chloroflexota, c: Dehalococcoidia, o: UBA6952, f: QGNO01, g: QGNO01 | 93.5 | iPHoP-RF, 95.40 |
| 10653 | 2412 | d: Bacteria, d: Actinomycetota, c: Actinomycetia, o: Actinomycetales, f: Actinomycetaceae, g: *Pauljensenia* | 92.6 | iPHoP-RF, 94.70 |
| 10955 | 2356 | d: Bacteria, d: Pseudomonadota, c: Gammaproteobacteria, o: Xanthomonadales, f: Rhodanobacteraceae, g: *Luteibacter* | 91.8 | iPHoP-RF, 94.10 |
| 11146 | 2325 | d: Bacteria, d: Actinomycetota, c: Actinomycetia, o: Actinomycetales, f: Micrococcaceae, g: Paenarthrobacter | 94.5 | iPHoP-RF, 96.10 |
| 11945 | 2199 | d: Bacteria, d: Bacteroidota, c: Bacteroidia, o: Bacteroidales, f: Muribaculaceae, g: JAGBWK01 | 90.3 | iPHoP-RF, 92.80 |
| 12142 | 2173 | d: Bacteria, d: Bacteroidota, c: Bacteroidia, o: Bacteroidales, f: Bacteroidaceae, g: *Prevotella* | 92.6 | iPHoP-RF, 94.70 |
| 12324 | 2148 | d: Bacteria, d: Pseudomonadota, c: Gammaproteobacteria, o: Enterobacterales, f: Enterobacteriaceae, g: *Serratia* | 92.6 | iPHoP-RF, 94.70 |

| | | | | |
|---|---|---|---|---|
| 12706 | 2102 | d: Bacteria, d: Actinomycetota, c: Actinomycetia, o: Mycobacteriales, f: Mycobacteriaceae, g: *Nocardia* | 95.3 | iPHoP-RF, 96.70 |
